# Hierarchy and heterogeneity in the recruitment of *B. subtilis* replication-restart proteins to stalled forks

**DOI:** 10.64898/2026.03.20.710068

**Authors:** Robert C. Raatz, Daniel R. Hammerl, Anna Kornyushenko, Peter L. Graumann

## Abstract

The restart of replication forks that have become stalled or disintegrated during the replication cycle is vital for all organisms, and in many bacterial species involves the conserved and essential DNA helicase PriA. PriA has been shown to physically interact with the C-terminus of SSB, which also binds to several other proteins involved in DNA repair and restart. It has been proposed that PriA is enriched at all replication forks in *Bacillus subtilis* via SSB interaction, such that it is instantly present to respond to a requirement for restart. Using single molecule tracking, we show that SSB and PriA are comprised of populations having very different diffusion constants, ruling out that PriA is co-migrating with fork-bound SSB. Indeed, PriA was only enriched at a subset of cells in exponentially growing cells, dependent on the C-terminus of SSB, but largely showed confined motion through the entire genome, searching for target sites in a transcription factor-like manner. Upon stalling of forks, SSB became highly enriched in all cells, suggesting a first line of response. PriA was also visibly enriched at forks following replication stress, in contrast to primosome proteins DnaD and DnaI, who showed only moderate changes in localization or in single molecule motion. PriA dwell times were affected by the lack of the SSB C-terminus, and also by the absence of RecG helicase, which is involved in recombination events. Heterogeneity of restart proteins at replication forks also extends to translesion DNA polymerases PolY1 and PolY2. Both proteins are low-abundant such that a considerable fraction of cells is devoid of any molecule. Our findings show that SSB accumulation is an initial response to replication stress, and that translesion synthesis and lesion skipping are less frequent events than fork remodelling.

## Introduction

Duplication of genomically encoded information is one of the core elements of cellular life. During DNA replication, the complex molecular machinery faces different obstacles on the chromosomes or plasmids and has to overcome these, and must therefore be able to efficiently deal with conflicts such as transcription-replication-collisions, when the replisome has head-on encounters with RNA polymerase (RNAP), or breaks in the sugar-phosphate backbone of DNA, or base modifications that stall the progression of DNA polymerases (Bainbridge *et al*, 2021; Gabbai & Marians, 2010; Michel & Sandler, 2017; Torres *et al*, 2025; Windgassen *et al*, 2018). Upon encountering obstacles, the replication machinery may have to a) be able to bypass a base-modification or lesion, e.g. by employing translesion (TLS) polymerases that can put any base opposite a lesion, b) skip the lesion on the leading strand and re-load the polymerase behind the lesion (leaving time to fix the lesion post-replication), or c) move the replication forks backwards to allow removing the block or fixing the lesion by DNA repair pathways, or even completely dis - and reassemble the entire machinery (Heller & Marians, 2006; Torres *et al*., 2025). Failure to restart replication is detrimental, and therefore dealing with the many possible scenarios of lesion/block bypass is of outmost importance for all kinds of cells. This is emphasized by the fact that a key player in replication restart in most bacteria, PriA, is either essential for viability (e.g. in *B. subtilis*), or its absence causes severe defects. *E. coli priA* null mutant cells exhibit very slow growth, poor viability, and filamentation (Grompone *et al*, 2004). Also several components of the machinery initiating replication at the origin of replication (*oriC*), called the primosome, are proposed to be essential for replication-restart.

PriA is a widely conserved 3′-5′-DNA-helicase, consisting of several domains, not all of which are essential for replication restart (Gabbai & Marians, 2010). The N-terminal OB-fold is referred to as the (DNA) 3′-binding domain (3′-BD) and is necessary for survival of DNA damage, in addition to the winged-helix-domain (WH) that connects the 3′-BD with the C-terminal helicase domain. N- and C-terminal domains act together, resulting in stronger binding to single-stranded DNA (ssDNA) than just the helicase domain alone (Matthews & Simmons, 2022). The substrate range of PriA includes D-loops and Y-shaped DNA structures, and more specifically the junction of single-stranded (ssDNA) and double-stranded DNA (dsDNA) at stalled forks; the protein also possesses high affinity for the 3′-OH end of the leading strand (Lee *et al*, 2021). DNA-binding of PriA is not limited to replication forks only, but to DNA junctions in a broader context. Deletion of either of those domains results in hypersensitivity to DNA damaging reagents, but cells are still able to grow. PriA has several interaction partners: it binds to PriB and PriC in *E. coli*, which in turn leads to loading of DNA helicase DnaB via the recruitment of helicase loader DnaC (Michel & Sandler, 2017). PriB and PriC are found in Gram-negative bacteria, but not in e.g. *bacillota.* The *B. subtilis* primosome likely comprises DnaB, DnaD and DnaI. During initiation of replication at *oriC*, DnaA recruits DnaD, which associates with DnaB (not to be confused with *E. coli* DNA helicase DnaB) to recruit the DnaI/DnaC complex, with DnaC being the *B. subtilis* DNA helicase (Winterhalter *et al*, 2023). DnaI actively loads DnaC onto the chromosome (Ioannou *et al*, 2006), initiating establishment of replication forks. PriA also interacts with primosome proteins during replication restart (Bruand *et al*, 2001).

A second, key binding partner is SSB. Note that *B. subtilis* possesses SsbA at the replication machinery, and SsbB during DNA uptake in cells grown to competence. We will refer to SsbA as SSB, to avoid confusion. Lecointe *et al*. identified an 8C-motif in PriA that resembles a zinc-finger-domain, whose loss leads to an abolishment of pas-dependent stimulation of ATPase activity, and loss of interaction with SSB (Lecointe *et al*, 2007). *Bacillus subtilis* PriA_Bs_ showed a preference for interaction with SSB that is bound to single-stranded DNA (ssDNA). Interestingly, an ectopically induced GFP-PriA fusion formed discrete foci in the cell, at replication forks, dependent on the (non-essential) C-terminus of SSB. The C-terminal acidic tip of SSB also interacts with proteins involved in homologous recombination (HR) that should be targeted to stalled replication forks, such as helicases RecQ and RecG. Because SSB tetramers tightly interact with PriA, it has been proposed that the visible accumulation of PriA at replication forks facilitates replication restart, due to a proximity effect. Little is known about the recruitment of other primosome components to the replication machinery.

TLS polymerase Pol IV in *E. coli* has been shown to act at sites away from replication forks, i.e. during base excision or nucleotide excision repair (Henrikus *et al*, 2018; Thrall *et al*, 2017), and is recruited to the replication machinery upon induction of replication stress. Conversely, for PolY1 from *B. subtilis*, the polymerase was reported to be a regular constituent of replication forks (Marrin *et al*, 2024). This would imply that *B. subtilis* replication forks are also set to rapidly deal with sites of DNA lesions/damage that can not be processed by the major polymerase PolC. Also different from *E. coli, B. subtilis* uses two different DNA polymerases at the lagging strand: error-prone DNA polymerase DnaE extends the RNA primers generated by primase DnaG, and rapidly hands over to PolC, which finishes the larger part of the Okazaki fragment. It is still unclear if DnaE could be employed for lesion bypass at the leading strand, in addition to the two TLS polymerases PolY1 and PolY2.

Because a large variety of proteins have been implicated in replication restart using genetics and biochemical tools, and many different temporal scenarios have been proposed, we wished to analyse restart in live cells by following changes in molecule dynamics at a single molecule level. Single molecule tracking enables to elucidate replication restart dynamics at single protein level. PriA is naturally only present at low levels in the cell, and in earlier studies, association with forks was tested using GFP-PriA induced in addition to PriA expressed from the original gene locus, possibly leading to overproduction that may affect localization. We therefore generated a strain encoding mNeonGreen-PriA under its native promoter using CRISPR integration. Treatment with HPUra (inhibition of PolC) or Mitomycin C (DNA crosslinks) was employed to understand the response of PriA to initiate replication restart. SMT revealed major differences in the dynamics of DnaC, SSB and PriA under induced restart conditions, which are not compatible with a simple proximity effect for PriA. SsbA showed major changes in replication fork-bound and freely diffusing fractions after application of HPUra or MMC. Even diffusion constants of the two fractions varied greatly, indicating major changes in binding partners. Conversely, PriA showed only small changes, and even release from replication forks during repair of MMC damage. Replicative helicase DnaC showed minor changes as expected, which reveal that changes seen for SsbA are specific rather than general. Our data show that replication restart involves major changes at the lagging strand, and little change with respect to the DNA helicase.

## Materials and Methods

### Bacterial strains and growth conditions

Bacterial strains are listed in table S1, and primers in table S2. *Escherichia coli* XL-1 Blue and BW25113 were used for cloning and all *Bacillus subtilis* strains in this study were derived from the BG214 wild type, Cells were grown in LB-Miller medium at 30°C while shaking at 200 rpm. After reaching exponential phase at an OD600 of 0.5 - 0.6 cells were treated with or without HPUra, or Mitomycin C, for 15 min at 37°C and 700 rpm in a thermoshaker (Eppendorf). When required the following antibiotics were added: chloramphenicol at 5 µg/mL; spectinomycin at 100 µg/mL. When required 50 µg/mL 6-(p-hydroxyphenylazo)-uracil (HPUra) was added. Mitomycin C was added to a final concentration of 50 ng/mL.

### Strain construction

DnaI, DnaC, DnaX, PolY1 and PolY2 were fluorescently labeled by inserting 500 bp of the 3′-end (lacking the stop codon) into pSG1164-mNG (Fiedler & Graumann, 2024) to generate a C-terminal mNeonGreen-tagged variant via single-crossover. DnaD was labelled by inserting 500 bp of the 5′-end (including the ribosome binding site) into pHJDS-mNG to generate a N-terminal mNeonGreen-tagged variant by integration of pHJDS-mNG in the original locus, thereby exchanging the promotor with a xylose-inducible promotor.

To visualize dynamics of Y-polymerases relative to replication, Y-polymerase mNeongreen fusions were combined with a DnaX-CFP. The entire *dnaX* gene was cloned into pSG1192-cfp (Feucht & Lewis, 2001) and was integrated into the *amyE*-site of BG214, selecting for Spec resistance. The resulting strain was transformed with chromosomal DNA of strains carrying *polY1-mNG* or *polY2-mNG* fusions, selecting for Cm resistance.

SSB was introduced as an ectopically tagged version with mNeonGreen by using pSG1193-mNG. The plasmids were generated with NEB HiFi DNA assembly master mix and transformation of chemically competent *E. coli* XL-1 Blue. *Bacillus* transformations were done by using MC-media as previously described (Fiedler & Graumann, 2024).

A fusion of *priA* to *mNeongreen* was carried out using a CRISPR-Cas9 approach previously described by Burby and Simmons (Burby & Simmons, 2017). The start of the *priA* gene lies within the *coaBC* orf; to ensure proper translation of *coaBC* we used a modified version of *mNeonGreen* such that *coaBC* stays intact, using the primers listed in table S2. The second codon of *mNeongreen* was deleted for this purpose. CRISPR-Cas9 generated fusion of *priA* to *mNeongreen* was verified by colony PCR. Modified cells were heat treated at 42°C and checked for loss of pPB105. Afterwards cells were once again checked for modification by PCR and sequencing of the PCR product. To test for levels of PriA expressed from the *amyE* locus, *m-NG-priA* (full length) was inserted into pSG1193, which was integrated into the *B. subtilis* chromosome via double crossover integration. In this construct, expression of *mNG-priA* is driven by *pxyl*, in addition to the *priA* gene at the original gene locus.

SSBΔ35 was also generated using CRISPR-Cas9. The last 35 codons were deleted using the primers mentioned in table S2 to generate a repair template missing these codons. The downstream *rpsR* gene was not modified. Strain generation as described above for *priA-mNeongreen*.

A *recG* deletion strain was constructed using long-flanking-homology PCR from the respective deletion strain of the *Bacillus* gene deletion collection (BKK15870, Kanamycin-based) (Koo *et al*, 2017). A PCR product carrying the resistance gene and 2 kb upstream and downstream of *recG* was generated using the primers stated in table S2, and the strain expressing mNG-PriA was transformed with this fragment, excluding unwanted recombination events away from the deletion cassette.

### Sensitivity assays

Cultures were grown until an OD_600_ of 0.5-0.6. A 100 µL aliquot of the cultures was OD-normalized and serially diluted 10 fold in a 96-well plate. 10 µL of each dilution was spotted onto LB-agar plates containing Mitomycin C in different amounts. Plates were incubated for 24 h at 30°C.

### Single molecule tracking

For single molecule tracking cells were grown in 10 mL LB-media with the corresponding antibiotic(s) at 30°C and 200 rpm until an optical density of 0.5-0.6. ensuing, a 100 µL sample was taken, the reagent (HP-Ura or MMC) applied and the sample incubated at 37°C, 700 rpm for another 15 min. The untreated conditions were treated likewise without drug administration. Cells were imaged at room temperature using, an inverted microscope Nikon Ti-E eclipse with a 100 x, A = 1.49, oil-immersion objective and 1.6 x optovar magnification. The central part of a 514 nm Laser (extended to 2 mm width) was used at 20 mW power for bleaching and excitation. The laser intensity was adjusted in accordance with the protein abundance to minimize bleaching time without applying unnecessary heat stress. Movies were processed using ImageJ (Hammerl & Graumann, 2024). In correlation to the amount of protein present, the first 100 to 600 frames of the aquired 3000 frames were cut off. Afterwards tracks were identified by U-track 2.0 (Jaqaman *et al*, 2008) and analysed with SMTracker 2.0 (Oviedo-Bocanegra *et al*, 2021).

### Western-blotting

To verify the full length of all mNeonGreen-tagged proteins the cultures were harvested at OD_600_ of 0.6 and lysed in TE-buffer with 10 mg/mL lysozyme at 37°C, 700rpm. The cell lysate was mixed with SDS-sample buffer and incubated at 60°C for an hour to ensure proper defolding of proteins. Blotting was done by using a nitrocellulosemembrane. For blocking and washing steps PBS and PBST buffer were used. Detection was done by using a 1:1000 dilution of anti-mNeonGreen-α-rabbit-IgG antibody for primary detection and a 1:10000 dilution of α-rabbit-IgG-HRP-conjugated antibody for secondary detection with HRP substrate solution (VWR).

## Results

### A markerless N-terminal fusion of PriA at the original gene locus reveals heterogeneity in fork association

Previously, PriA has been shown to retain function when it is fused to a fluorescent protein at its N-terminus (Lecointe *et al*., 2007). The protein was shown to be present at all replication forks in exponentially growing cells, via binding to the C-terminus of SSB. However, the fusion was expressed under the control of the xylose promoter, from the ectopic *amyE* gene locus. Because PriA is present in low amounts (about 50 copies) in *B. subtilis* cells (Polard *et al*, 2002), even low expression of mRNA from a second gene locus will considerably increase protein copy number, with higher than normal protein levels possibly influencing the localization of the protein. We therefore generated a construct, using CRISPR technology, in which mNeongreen is fused to the *priA* gene at the 5’-end, such that an mNeonGreen-PriA (mNG-PriA) fusion is expressed from its original gene locus, i.e. under control of its original promoter, entirely retaining operon structure (Fig. 1C). Western blot analysis showed detectable expression of full-length mNG-PriA (Fig. 1A, note that lanes 3 and 4 contain higher protein loads), but much lower levels compared to DnaI, DnaC or DnaX (Fig. 1A, compare lane 7 with 11 to 13). When mNG-PriA expression is driven by pxyl at the original gene locus (Fig. S1, lane “mNG-PriA 0.5%”), cells contain considerably more protein compared with the CRISPR/Cas fusion with the original promoter (Fig. S1, “P”). Thus, prior studies using such a construct were likely prone to producing artefacts due to much higher amounts of PriA compared with physiological levels. Cells carrying the mNG-priA fusion showed survival to growth in plates containing Mitomycin C comparable to wild type cells (Fig. S2), showing that the fusion protein can functionally replace the wild type protein also in response to replication stress.

**Figure 1.**
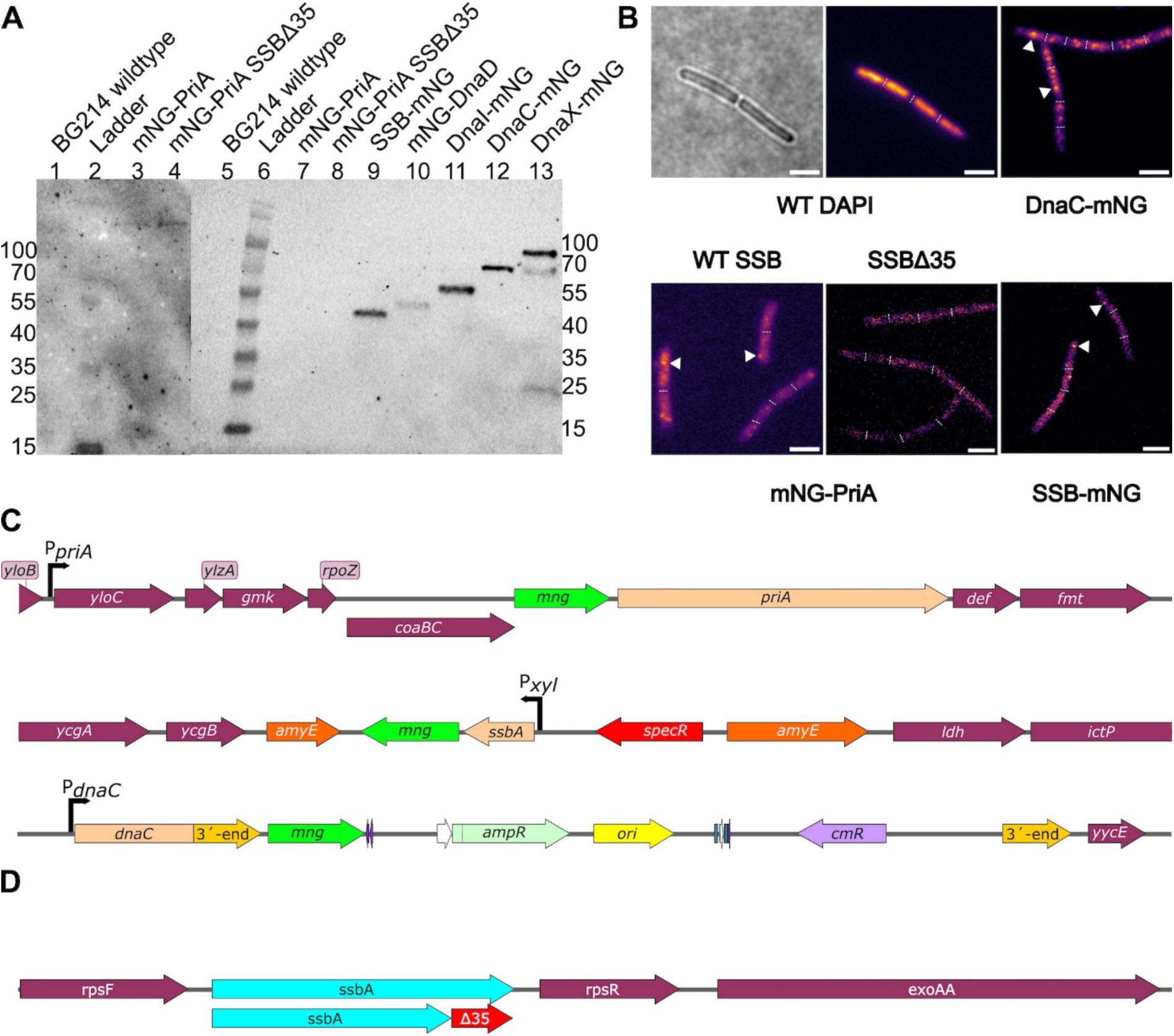
A) Western blot using anti-mNeonGreen antibodies. Same amounts of cell extract were loaded into each lane, except for lanes 3 and 4 containing more protein load from cell extract. SSB-mNG in lane 9 and mNG-DnaD are expressed using 0.05% xylose. B) Epifluorescence microscopy, “DAPI” DNA stain. Triangles in images showing mNG fluorescence indicate fluorescent foci. White bars 3 µm. C-D) Genetic environment of fusions. *mNG-priA* (C) and truncated *ssbA* (D) were generated using direct insertion/deletion via CRISPR/Cas technology.

Epifluorescence microscopy showed that mNG-PriA formed foci in some cells, but only in a minority (Fig. 1B, Fig. 2). In most cells, fluorescence showed a nucleoid-like pattern, suggesting that the protein is largely associated with sites throughout the entire genome, while only 29% of cells showed a clear accumulation of PriA at the replication forks. Visible recruitment to forks is indeed mediated by the C-terminus of SSB, because no fluorescent foci were detectable in cells expressing “SSB-delta35” (Fig. 1C, Fig. 2), supporting findings by Lecointe et al. (Lecointe *et al*., 2007). Thus, postulated enrichment of PriA at the forks to facilitate restart is only true for a sub-fraction of an exponentially growing population, suggesting that largely, PriA finds stalled replication forks by a diffusion-capture mechanism, or by other means. We will return to this idea in a later section.

**Figure 2:**
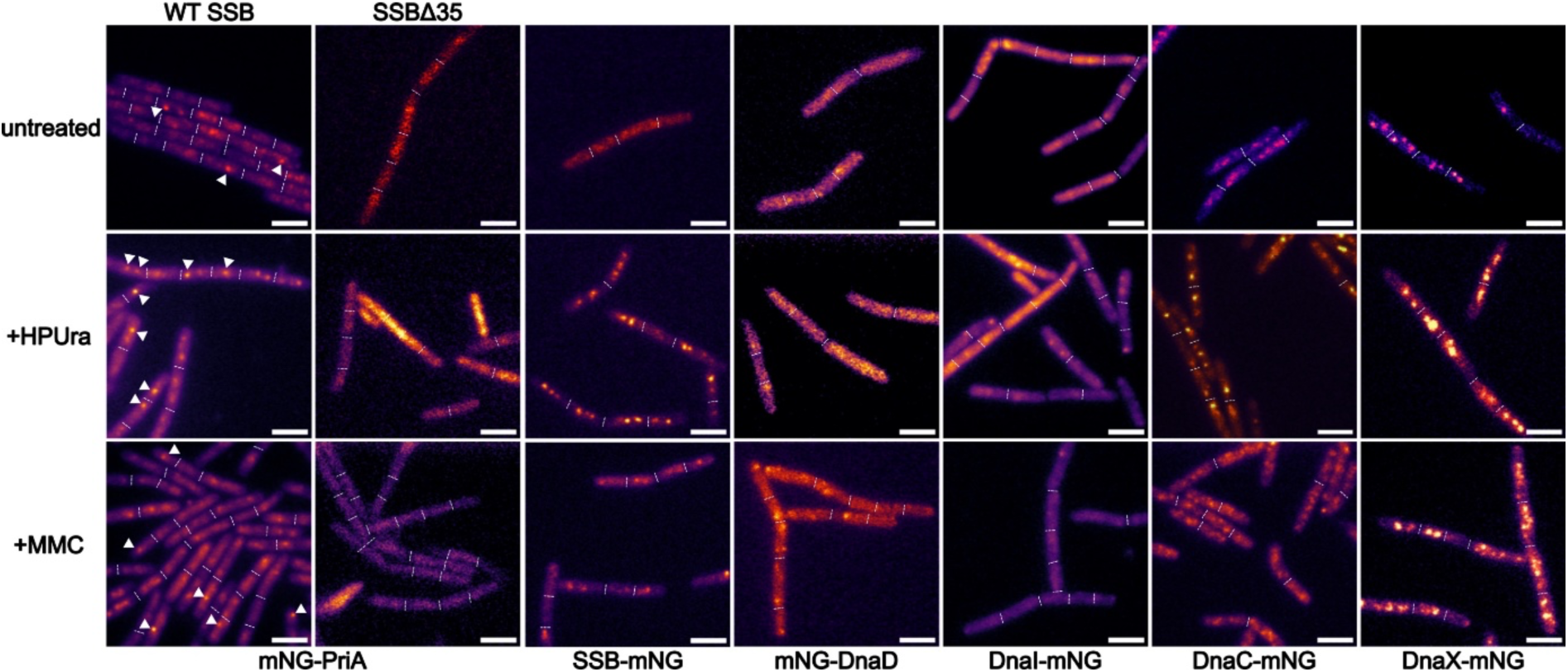
Epifluorescence imaging of PriA, DnaC and SsbA fusion proteins with mNeongreen (A). PriA is under control of its native promoter and shows distinct foci. Fluorescence signal increases after HPUra (215 µM) as well as Mitomycin C (MMC, 50 ng/mL) treatment. White triangles indicate mNG-PriA foci. Dashed lines indicate ends of cells. Scale bars 3 µm.

We also generated a strain in which a DnaC-mNeonGreen fusion is expressed from the original gene locus using single crossover integration of a pSG1164 derivative. DnaC is a monocistronic gene (Fig. 1C), so no downstream gene is affected by plasmid integration. Like the CRISPR-mNG-PriA strain, cells expressing DnaC-mNG as the only form of the helicase grew with a similar doubling time as wild type cells (38 ± 3 min), showing that the fusions can functionally replace the wild type proteins. As expected from cells growing with overlapping rounds of replication like in our case (LB medium, 30°C), a large majority of cells (>95%) contained DnaC-mNG foci, similar to DnaX-mNG (Fig. 1B), which was constructed similarly to the *dnaC-mNG* strain. These finding verify that PriA is only recruited to a subset of replication forks by SSB, revealing a clear heterogeneity in fork composition within an exponentially growing population.

### DnaD and DnaI are present as preassemblies in a subset of cells

We aimed at comparing the localization and dynamics of PriA with that of other replication restart proteins, and especially with SSB, because of the physical interaction between the two proteins. A C-terminal SSB fluorescent protein fusion has been shown to be able to functionally replace the wild type protein (Dubiel *et al*, 2020). However, the C-terminus is required for binding to PriA or other repair proteins (Lecointe *et al*., 2007), so a C-terminal fusion potentially interferes with protein/protein interactions, which is especially important during conditions of replication stress. To circumvent this caveat, we expressed the SSB-mNeonGreen fusion protein from the *amyE* locus using low xylose induction (Fig. 1A). Because SSB is a high-abundant protein (between 2000 molecules per cell in minimal medium to 14000 in rich medium molecules per cell (Dubiel *et al*., 2020)), low, merodiploid expression of SSB-mNG will ensure that SSB tetramers only contain a single fusion protein and very rarely two. This increases a) the likelihood of that SSB-mNG-containing tetramers behave similar to wild type tetramers, and b) that a majority of C-termini (also in tetramers with an incorporated SSB-mNG) remain free for binding to PriA or other proteins.

Finally, we generated a C-terminal mNG fusion to DnaI, and an N-terminal fusion (driven by pxyl) to DnaD. Both strains grew with a doubling time of 39 ± 3 min, or 42 ± 4 min minutes, respectively, in rich medium at 30°C, compared with 38 min for cells lacking any fusion construct, showing that functionality was retained during exponential growth. For DnaD, it is possible that the protein was overproduced, and that overproduction may mask a reduced, partial functionality. Fig. 1A shows that all fusions were expressed as full-length proteins. Visualization of fusion proteins by epifluorescence revealed that DnaD and DnaI formed foci in a minority of cells (DnaD: 37%; DnaI: 19%, Fig. 2), similar to PriA, but distinct from DnaC or DnaX that showed discrete foci in >90% of the cells. Thus, for three restart proteins, association with replication forks shows heterogeneity between cells.

### Replication stress leads to strong changes in the intensity of foci of SSB

We generated two different types of externally induced stress for the replication forks, (reversible) inhibition of PolC by HPUra, and stalling of entire forks when DnaC and polymerase activities are compromised by Mitomycin C (MMC), which generated base adducts and crosslinks between the two DNA strands. Interestingly, mNG-PriA now became visible at >82% of cells as one or more fluorescent foci (Fig. 2), indicating that the protein now finds a majority of forks and accumulates there. Intriguingly the intensity of SSB-mNG foci strongly increased (Fig. 2), in spite of maintained (low) levels of SSB-mNG expression by pxyl. This finding indicates that there is a highly increased amount of substrate binding sites at stalled forks, i.e. a highly increased degree of ssDNA, suggesting continued activity of the helicase, running far ahead of stalled polymerases. Less dramatically than SSB, but clearly visible, the intensity of DnaC and DnaX foci was enhanced (Fig. 2), indicating an increased number of proteins at replication forks upon PolC stalling or MMC-induced fork stalling/collapse (of note, all images were taken at an exposure of 2 seconds). This would be compatible with some forks reloading DnaC, or an entire new replication machinery. However, as will become clear later on, this is likely not the case, because single molecule dynamics of DnaC and DnaX do not change markedly following replication stress (see below). Different from PriA, DnaD did not show a significant increase in focus formation following HPUra treatment, from 37% in non-perturbed cells, to to 41%, and an even decreased percentage of 21% after MMC treatment. These data must be viewed with the caveat that cells expressing DnaI-mNG showed a reduced capacity to grow on MMC-containing plates, but retained considerable viability, while mNG-DnaD expressing cells were strongly impaired in growth on the DNA-damaging agent (Fig. S2). Thus, changes in DnaD dynamics may be flawed in cells witnessing (continuous) replication stress.

DnaI-mNG showed a noticeable increase from 19% to 29% in cells with stalled PolC molecules, and following MMC treatment to 27%. In contrast to PriA, DnaD and DnaI frequently showed patch-like accumulation rather than discrete foci, wherefore quantification was somewhat subjective. Nevertheless, these visual data show that PriA accumulates at a vast majority of stalled replication forks, while DnaI and DnaD get recruited only to some forks during replication stress, and to different extents after MMC treatment or stalling of PolC.

Because quantification of epifluorescence images requires very careful calibration controls, we moved to single molecule tracking, which is inherently quantitative.

### PriA shows strikingly different mobility compared with SSB

All fusions were tracked with a 20 ms integration time, which allows to efficiently monitor freely diffusing cytosolic proteins. Fig. 3A shows the probability density function of jump distances (JD), obtained from squared displacement analyses (SQD). Assuming two populations of molecules with different mobility could well represent the observed distribution of jump distances, the darker lines in the JD plots in Fig. 3A are the sums of the two Rayleigh fits. For SSB and DnaC, it appears reasonable to assume a) freely diffusing molecules (with a diffusion constant D of 0.53 µm^2^ s^-1^ for SSB and 0.35 µm^2^ s^-1^ for DnaC) and b) molecules tightly bound at replication forks (D = 0.08 or 0.04 µm^2^ s^-1^, respectively), which is plausible because neither protein has been proposed to non-specifically bind to the chromosome, but to be directly recruited via protein/protein interactions or due to the presence of single stranded DNA behind the helicase. Indeed, using two Rayleigh fits can well explain the observed distribution of the probability density of jump distances (Fig. 3A-C). Fig. 3D shows a summary of squared displacement analyses (SQD).

**Fig. 3.**
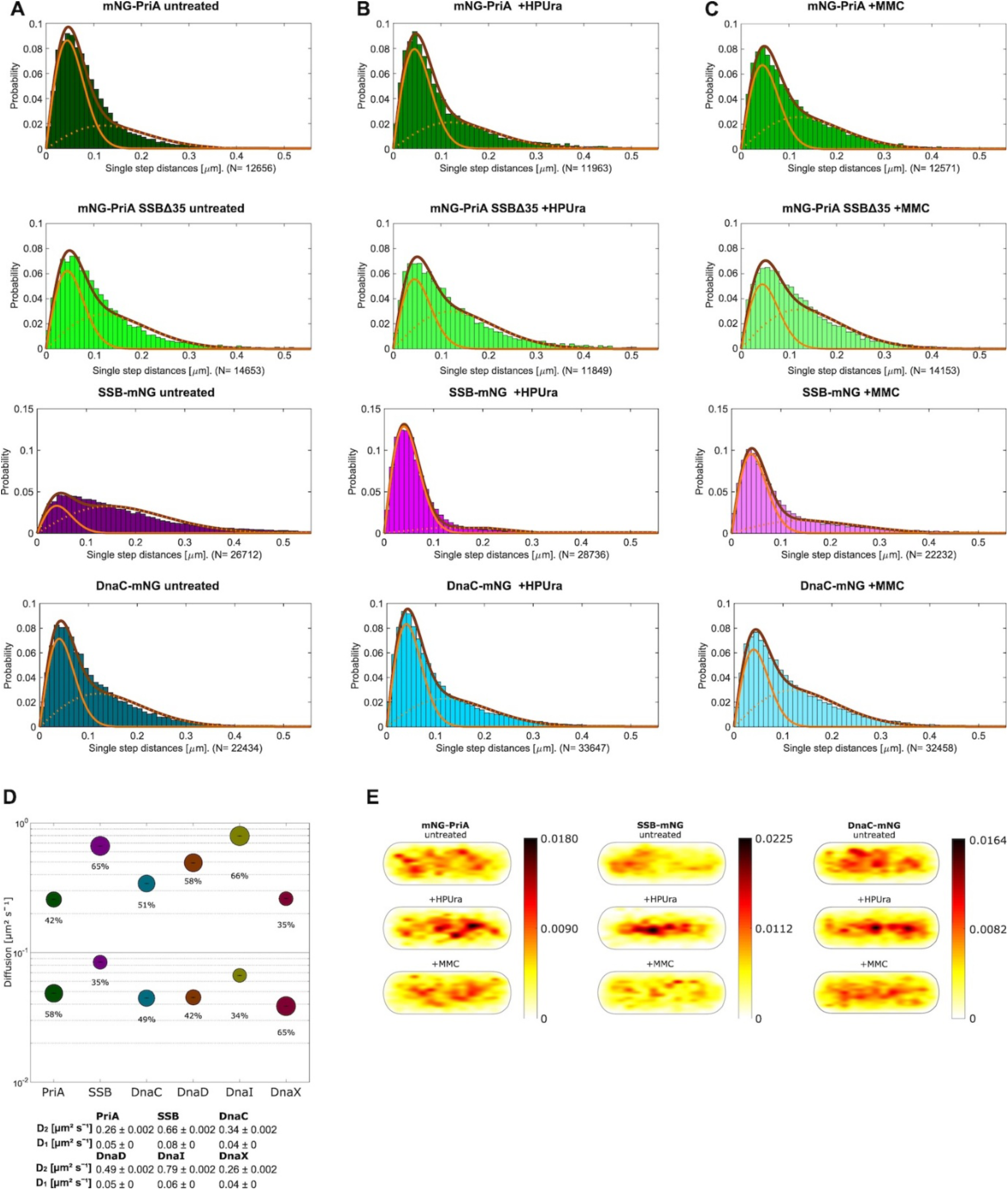
A-C) Jump distance plots based on Squared displacement analyses. Solid thin lines slow-mobile population, dashed lines fast-mobile populations, dark solid lines double fit. D) Bubble plots showing fraction sizes (size of the bubbles) and average diffusion constants on the Y-axis. Table of diffusion constants with mean fitting errors. E) Heat maps showing probability of localization in a medium-sized cell of 3 x 1 µm. Scale of most likely (dark) and most unlikely (yellow) localization besides the panels.

As expected, SSB being the smallest protein shows the highest diffusion constant for the fast-moving, freely diffusive population, and PriA being the largest displays the lowest diffusion constant. For PriA, 45% are in the freely diffusive state, while 55% are engaged in a low mobility state, likely engaged in a large complex and/or bound to DNA, while only 36% of SSB molecules are in a replication fork-engaged mode, and 64% are freely mobile (Fig. 3B). Assuming a conservative number of 2000 SSB molecules per cell, this would correspond to 180 SSB tetramers being present at two to four replication forks per cell, sufficient to coat more than 6000 nucleotides. Because SSB is not known to bind to dsDNA, and thereby find ssDNA regions via a scanning mode, an excess of free molecules supports rapid exchange of SSB at the lagging strand loops.

For DnaC, about half of the molecules were fork-bound, and half were freely diffusive. Interestingly, the static fractions also displayed very different mobilities: static PriA or DnaC molecules showed similar diffusion constants (0.05 or 0.04 µm^2^ s^-1^), while that of SSB was much higher (0.08 µm^2^ s^-1^) that that of PriA and DnaC. This may be based on the higher mobility of SsbA covering more flexible ssDNA regions at the lagging strand, detached from the major helicase/clamp loader/DNA polymerase complex. If a majority of PriA molecules were stably attached to SSB tetramers, or freely diffusive, we would expect similar diffusion constants for the faction of replication fork-bound SSB/PriA molecules. Thus, in unperturbed cells, static PriA molecules show DnaC-like static motion, incompatible with a major degree of recruitment to lagging strand-bound SSB molecules.

### SSB displays major replication stress-induced changes in mobility

Next, we analysed changes in protein dynamics resulting from replication stress. Fig. 4 shows that DnaC became slightly but considerably more static when PolC was inhibited by HPUra, and slightly more diffusive upon MMC treatment (blue panel). The static fraction increased by 17%, from 47 to 55% after treatment with HPUra, a change lying above a range of 10% difference, which we view as biological noise. Interestingly, SSB showed a major change in its dynamics upon addition of HPUra or of MMC; in both conditions, the molecules became drastically more static, from 22% static motion to 84% (Fig. 4, purple panel). Note that the values are different from Fig. 3B, because we have used a fitting procedure that finds the most suitable average diffusion constant for populations across all three conditions. This leads to changes only occurring in the size of populations, rather than in different average diffusion constants and different population sizes. This procedure allows for an easier assessment of relative changes in mobility. The findings suggest that inhibiting the progression of DNA polymerases does not strongly affect DnaC dynamics at the forks (some forks where lesions are bypassed by loading a new DnaC hexamer upstream of the lesion), but leads to massive loading of SSB, likely due to continued production of ssDNA by DnaC. These data support the idea that helicase continues unwinding of the DNA double helix, getting uncoupled from polymerase (in)activity. Interestingly, PriA dynamics did not follow the pattern of change seen for SSB. PriA became slightly more mobile after induction of replication stress by HPUra, and considerably less static after addition of MMC (Fig. 4, dark green panel). These experiments show that PriA and SSB show strikingly different motion at a single molecule level with regards to stalled forks.

**Fig. 4.**
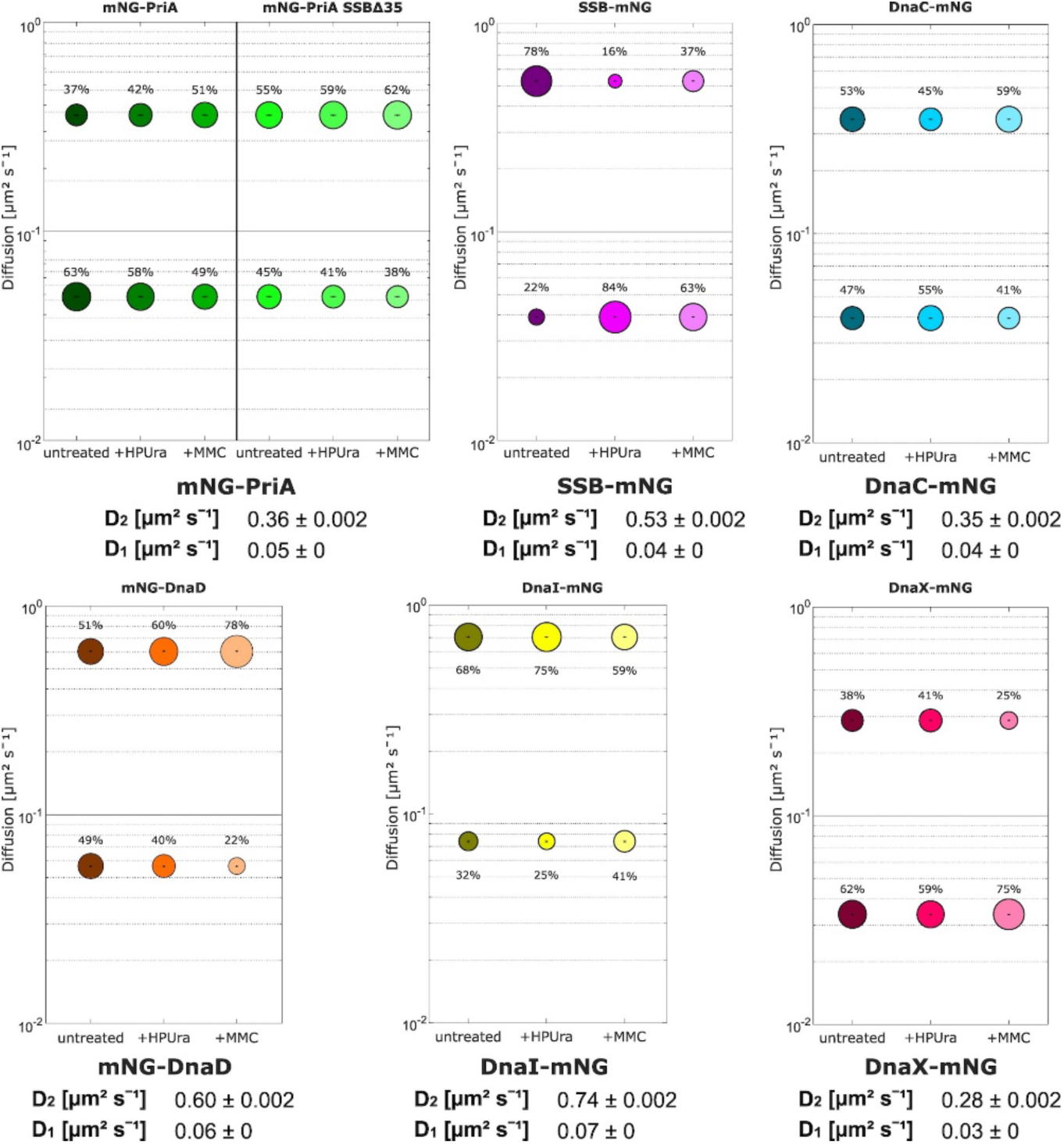
Bubble plots showing changes in protein mobility following replication stress. Diffusion constants are also stated underneath the panels.

We also analysed all molecular motion by projecting all tracks into a single, average-size cell. Heat maps shown in Fig. 3E show a preference of PriA, SSB and DnaC molecules for the central space of the cells, in agreement with their engagement with the chromosome. However, after stalling of DNA polymerases, all three proteins clearly accumulate stronger with central areas of the cell, in support of their higher association with replication forks on the nucleoid. This is especially pronounced for SSB, which shows the strongest changes in its single molecule dynamics, revealing that SSB molecules indeed become static in the areas containing replication forks, and show the strongest relocalization of all proteins analysed (heat maps for all proteins see Fig. S3).

The mobility of DnaX did not change considerable after stalling of PolC, suggesting that inhibition of the DNA polymerase also halts the procession of the clamp loader but does not lead to disassembly of the complex (Fig. 4, red panel). DnaX became more statically engaged on the chromosome (i.e. at the forks). This could reflect that MMC-induced lesions can stall the machinery and be bypassed by reloading a new machinery upstream of the lesion. Different to PriA, DnaI became more mobile following HPUra treatment, but more static after addition of MMC (Fig. 4, yellow panel). DnaD showed less static engagement following stalling of PolC and especially after MMC treatment (Fig. 4, orange panel) – keeping in mind that the mNG-DnaD fusion only poorly supported viability in cells growing under continuous replication stress (Fig. S2). These finding suggest that restart routes do not comprise strong changes DnaI dynamics, and possibly also DnaD motion, except for MMC-induced stress for DnaI.

In agreement with the idea that some PriA molecules are associated with replication forks, even under non-perturbed conditions, the fraction of slow-mobile/static PriA was considerably lower in a strain expressing SSB lacking the C-terminal interaction hub (Fig. 4, light green panel). However, there was still a large number of molecules showing motion that can only be explained through binding to a much larger complex, suggesting that some PriA molecules are still able to interact with replication forks.

### Cas9-like scanning of nucleoids by PriA

We have shown that the localization of mNG-PriA in many cells can be described as a nucleoid stain (Fig. 1). The presence of a non-specific DNA binding domain in PriA supports the idea that there is a DNA-bound fraction that moves through the nucleoid(s), via non-specific interactions with DNA, in addition to a freely diffusing (non-DNA bound) population and a fraction that is stably bound at the replication machinery.

To further test this idea, we generated Z-stacks of 500 ms time series (25 frames), called “sum” images. Fig. 5A shows examples of sum images, in which the signals from PriA molecules (mobile and static) add up as a clear nucleoid stain. The absence of clear foci recapitulates that a) most cells do not contain PriA at replication forks (in this setting during a 500 ms time interval), and b) that a majority of PriA molecules is associated with the nucleoid(s). SSB and DnaC showed clear foci or whole-cell fluorescence, although a nucleoid-like pattern was also observed in some DnaC-mNG cells (Fig. 5A). We also generated confinement maps for individual cells expressing mNG-PriA. Tracks confined to an area of 120 µm (three times the localization error) for at least 8 frames are shown in red, and localize to central areas containing the nucleoids, while most mobile tracks are also found in a pattern comparable to that of a nucleoid stain (Fig. 5B). Green tracks represent molecules changing between confined and non-confined motion. Overall, 60% of PriA tracks were freely diffusive, while 40% showed confined motion, i.e. stayed within a radius of 120 nm for at least 8 time intervals (table S3). The fact that in about 40% of the cells, only non-confined tracks are present, and in the 60% of cells all types of mobilities (Fig. 5B), suggests that PriA frequently changes between chromosome binding (be it by directly interacting with DNA or with DNA-bound molecules) and chromosome-scanning mode.

**Figure 5.**
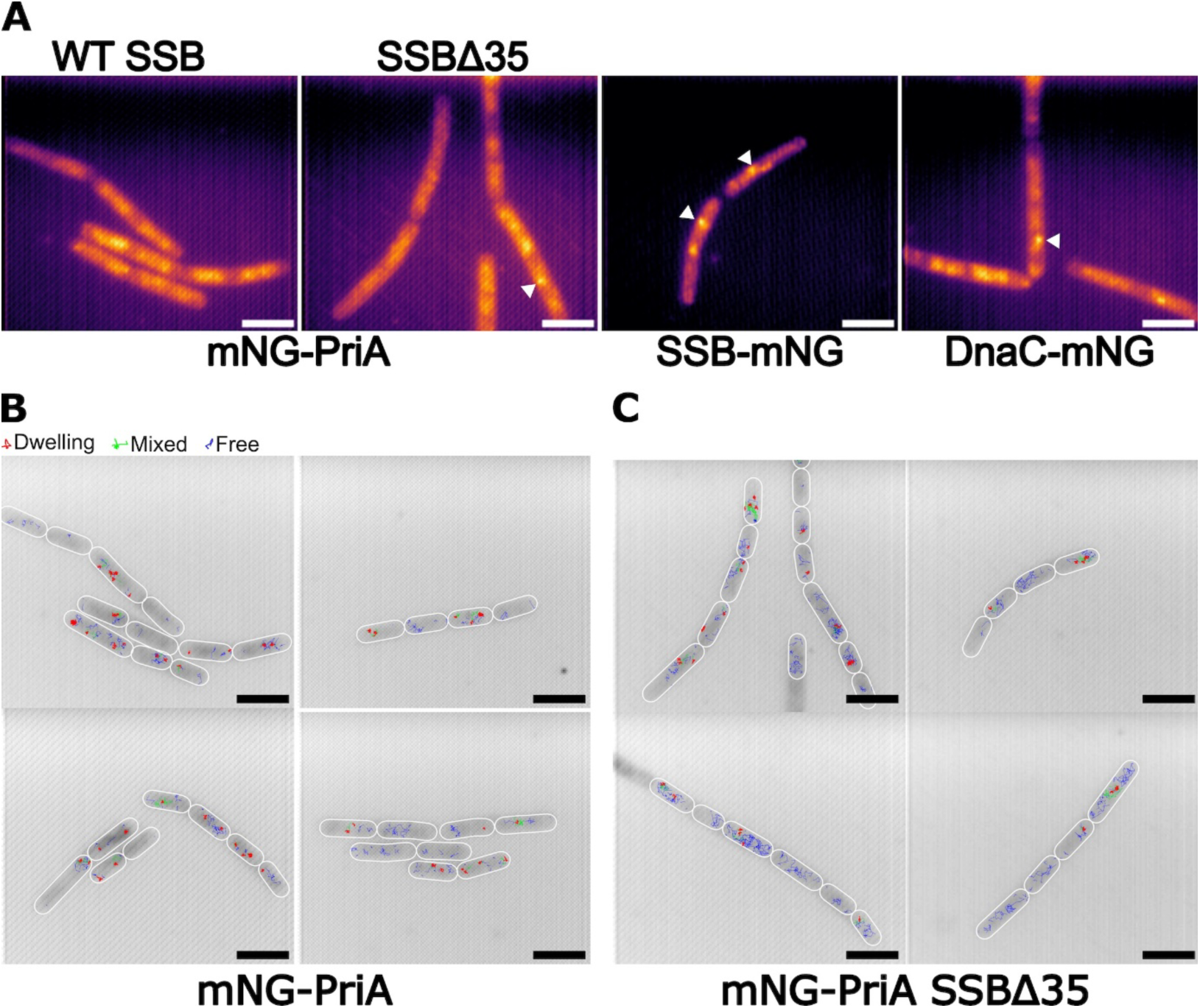
A) sum images of mNG-PriA in wild type cells and in cells lacking the C-terminus of SSB. Triangles point out discrete foci arising from long-dwelling molecules. Scale bars 3 µm. B-C) confinement maps of representative fields of cells; B) wild type cells, C) cells lacking the last 35 amino acids of SSB. Scale bars 3 µm.

Predominant motion on the nucleoids and long binding events have been described for bacterial transcription factors (Stracy *et al*, 2021), for Cas9 target search in *Lactococcis lactis* (Martens *et al*, 2019) and for a plasmid-encoded Type IV CRISPR/Cas system in *Pseudomonas oleovorans* that targets a single site on the chromosome to silence transcription (Guo *et al*, 2022). Our observations indicate that PriA uses a similar mode of 1D motion along DNA strands and 3D hopping between strands to find preferred binding sites.

We further analysed if PriA shows extended dwell events in cells lacking the C-terminus of SSB. Fig. 5C shows that PriA still showed dwell events, but overall, 25% of PriA molecules showed confined motion, rather than 40% as in wild type cells, and 75% moved freely (table S3). Thus, while visible accumulation of PriA at stalled replication forks is affected by the lack of the SSB interaction domain, PriA retains a nucleoid-associated scanning mode including extended arrests in the absence of the C-terminus of SSB.

### PriA dynamics are strongly affected by the lack of the C-terminus of SsbA and in the absence of RecG

As shown above, PriA shows strong association with the *B. subtilis* nucleoids, suggesting the existence of three populations with distinct average mobility. Using the Bayesian information criterion implemented in the analyses software, SMTracker 2.0, suggests that using a three-population fit for the JD analyses of PriA does not represent overfitting of data. We also compared fitting of SQD data using two or three Rayleigh fits, with the latter resulting in a more accurate description of the observed distribution of displacements (suppl. Fig. S4). This analysis suggests that PriA shows three distinct modes of motion: besides free diffusion of a small fraction of PriA molecules (about 17%), there is a major, medium-mobile population scanning the chromosome (64%), and about 19% of molecules statically engaged in DNA/replication fork binding (Fig. 6). Based on the assumed existence of three populations, stalling of DNA polymerase results in increased binding to replication forks (from 19 to 43%), at the expense of the nucleoid-scanning molecules (decreasing from 64 to 36%) and an increasing freely mobile population (Fig. 6, middle panel). Given that the number of foci of mNG-PriA increases after HPUra treatment (Fig. 2), these data appear to better explain the single molecule dynamics of PriA. In response to MMC treatment, PriA also showed increased free diffusion as well as increased static motion, at the expense of the medium-mobile population (Fig. 6, right panel). Lack of the C-terminus of SSB led to a decrease in the fraction of static PriA molecules under all conditions (Fig. 6, light green bubbles), but did not abolish the static fraction, which remained as a considerable proportion of 12 to 26% of molecules.

**Figure 6.**
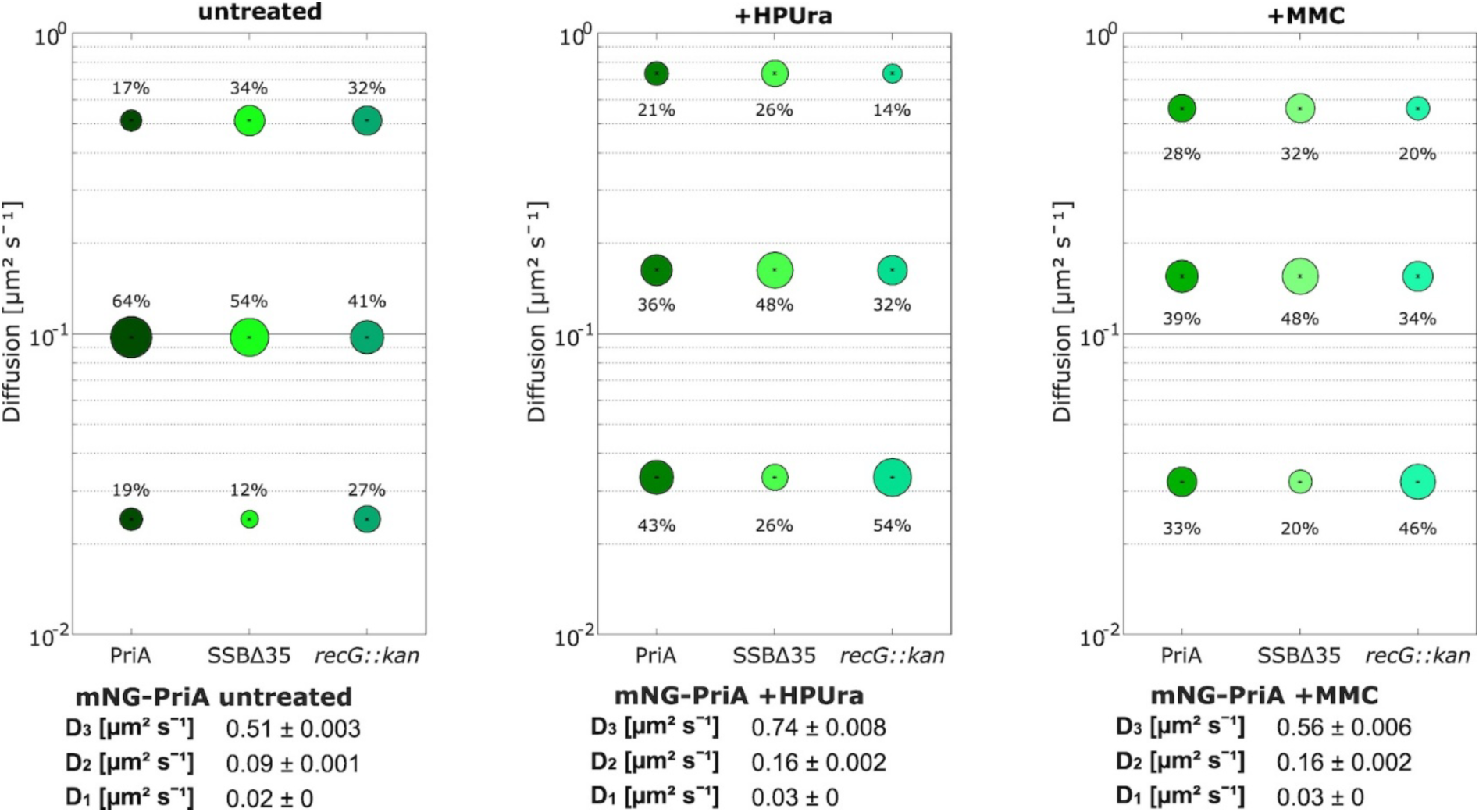
Bubble plots showing fraction sizes and their diffusion constants. Simultaneous fits for all three genetic backgrounds were employed.

Helicases RecG and RecQ are involved in DNA repair via HR, and have also been reported to be recruited to replication forks via binding to the C-terminus of SSB (Lecointe *et al*., 2007). RecG has additionally been implemented in replication restart based on genetic evidence in *E. coli* (Gabbai & Marians, 2010; Rudolph *et al*, 2009). We wished to investigate if the absence of RecG has an effect on the mobility of PriA molecules, and therefore deleted the *recG* gene in the mNG-PriA strain. Fig. 6, left panel, shows that contrary to a reduction of the static PriA fraction in cells lacking the C-terminus of SSB, lack of RecG strongly increased this fraction, from 19 to 27% (medium-dark green bubbles). Note that again, numbers are somewhat different because wild type and mutant cells are compared assuming the same diffusion constants for PriA populations. Increased static population was accompanied by a decreased medium-mobile population. Thus, PriA remains more strongly engaged in chromosome-binding in the absence of RecG, which is retained during replication stress via HPUra (Fig. 6 middle panel) or MMC (right panel). These finding reveal a strong influence of RecG on the dynamics of PriA, indicating a strong connection between PriA activity and proteins involved in homologous recombination. RecG has been suggested to play an important role in fork remodelling, e.g. by the reversal of stalled forks (Torres *et al*, 2021); if RecG acts downstream of PriA, its absence would increase engagement of PriA with stalled forks. Alternatively, cells lacking RecG may contain an increased number of D-loops, and thus more substrates for PriA to bind to.

We calculated average dwell times from cumulative probability density functions of dwell events, taking into account dwell events of at least 8 frames. Fitting of decay curves to the data suggests the existence of two distinct populations having different average dwell length (Fig. S5). In wild type cells, PriA showed an average dwell time of 0.324 ± 0.013 s, in cells lacking the C-terminus of SSB reduced dwelling of 0.296 ± 0.014 s, and in delta *recG* cells increased dwell time of 0.340 ± 0.013 s. Considering the two different fraction, in wild type cells, 48.6 ± 4.6% showed shorter dwell events of 200 ms, while 51.4 ± 4.6% had extended average dwell times of 400 ms. In the absence of RecG, 59% of cells had a dwell time of 220 ms, and 41% showed dwelling events of 480 ms. Thus, the increase in the static fraction of PriA in *recG* mutant cells is accompanied by measurably increased dwell events, in a manner opposite to the lack of the protein-interaction terminus of SSB.

### Low abundance of PolY1 and PolY2 generates heterogeneity in molecule availability for replication forks between cells

*B. subtilis* PolY1 and PolY2 belong to the Y-family or error-prone TLS polymerases, which can polymerize through positions containing damaged bases that can not be processed by PolC. PolY1 has been shown to lead to an increased mutation rate (Duigou *et al*, 2004), in particular for genes on the lagging strand (Million-Weaver *et al*, 2015), by contributing to elevated mutagenesis of genes transcribed in a head-on orientation with replication (Million-Weaver *et al*., 2015). However, overall mutation rate is similar between wild type and *polY1* mutant cells (Carvajal-Garcia *et al*, 2023; Lenhart *et al*, 2012), possibly due to redundancy with other DNA polymerases. PolY2 has been implicated in UV-induced mutagenesis, and ist transcription is upregulated during the SOS response. PolY1 has been described as a constituent of the DNA replication forks in *B. subtilis*, via binding to the beta-clamp, DnaN (Marrin *et al*., 2024). This is different from *E. coli* TLS polymerases, which are recruited to replication forks that are stalled, via interaction with the clamp, but are absent from forks (i.e. not enriched) during unperturbed cell growth. In general, TLS polymerases are also implicated in DNA repair via NER and BER, because a) they interact with PolA (Duigou *et al*, 2005), which fills gaps in the DNA generated during NER and BER, and b) Pol IV in *E. coli* shows binding events on the chromosome away from replication forks during unperturbed growth (Thrall *et al*., 2017).

The finding that PolY1 is accumulated at non-perturbed *B. subtilis* replication forks suggests that translesion synthesis can occur rapidly, by exchanging PolC for PolY1 associated at the beta clamp. We wished to investigate the single molecule motion of PolY1 and of PolY2 in exponentially growing *B. subtilis* cells, relative to replication forks. Both fusions were generated at the original gene locus, leaving intact their transcriptional regulation and expression levels.

SMT of PolY1 showed a JD distribution that could be fitted with a three-population model without overfitting of data, according to the SMTracker software. It is reasonable to believe this in true *in vivo*, as DNA polymerases have a general DNA-binding ability (albeit much lower than in conjunction with the beta clamp), and would therefore move through the chromosome analogous to a transcription factor, besides tightly bound or freely diffusive mode. Apparent diffusion analyses suggests that PolY1 shows three distinct populations, with diffusion constants of 0.056 µm^2^ s^-1^, 0.31 µm^2^ s^-1^, or 1.06 µm^2^ s^-1^, compatible with a large, freely diffusive fraction (48%, Fig. 7A), an also quite large medium-mobile (likely chromosome-scanning) fraction of 38%, and a much smaller static (tightly chromosome-engaged) fraction of 14%. Interestingly, we found tracks for PolY1 (during the 60 seconds experiments) in only 84% of all cells. Fig. 7A shows that in contrast to some cells containing a considerable number of tracks (Fig. 7B), some cells did not show any PolY1 tracks. While replication forks – visualized via DnaX-CFP – did show association with dwelling PolY1 molecules, most forks were devoid of any PolY1 signal. These data show that the number of PolY1 molecules per cell is generally so low that there are not enough molecules present for each replication forks.

**Fig. 7.**
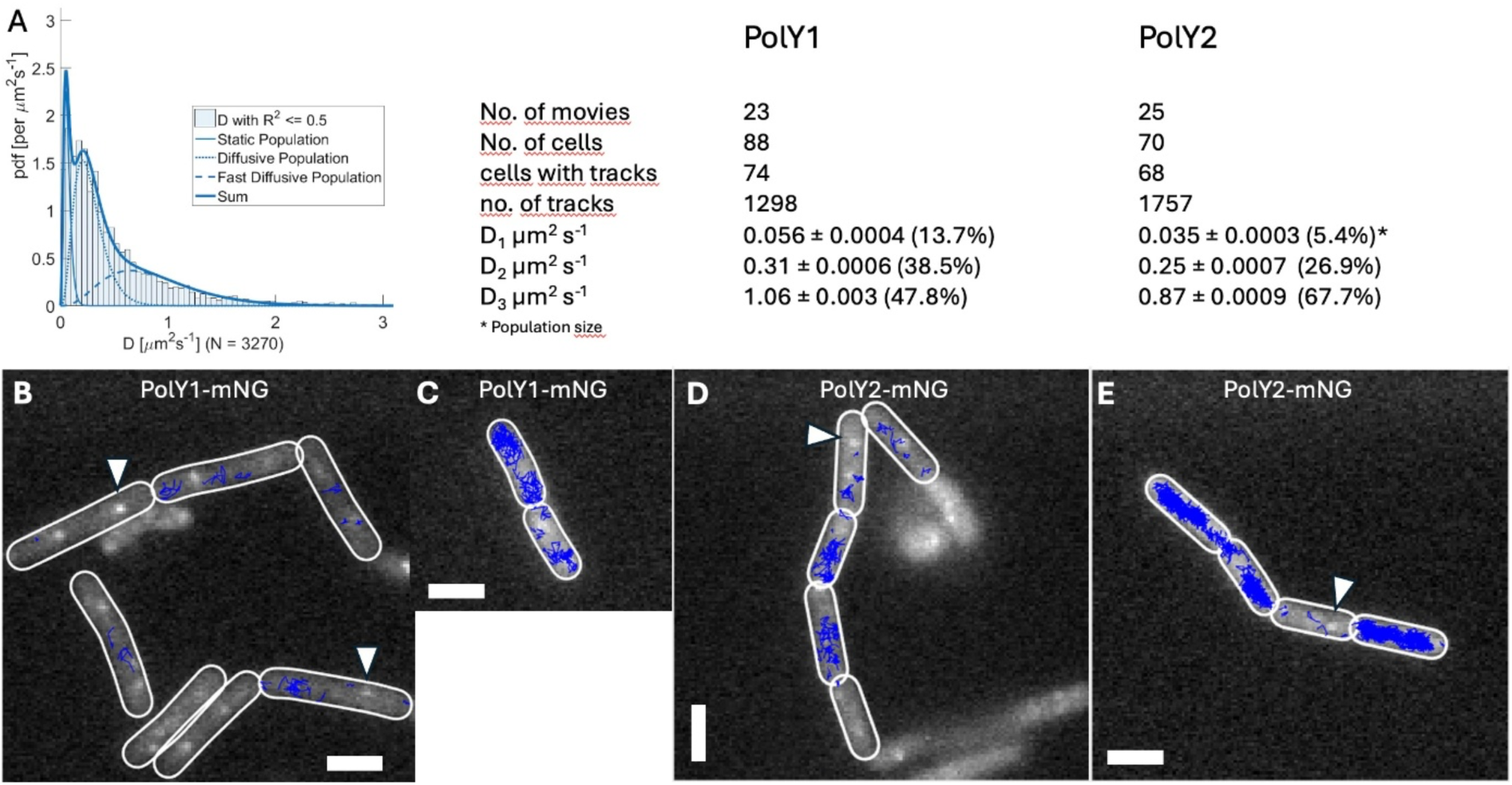
A) Apparent diffusion histogram for PolY1-mNG during exponential growth. B-E) overlay of fluorescence from DnaX-CFP and tracks of PolY1-mNG (B-C) or PolY2-mNG (D-E). Scale bars 2 µm.

Surprisingly, we found a higher number of tracks per cell for PolY2, which was earlier supposed to be only synthesized during the SOS response (Sung *et al*, 2003). We found 97% of cells containing PolY2-mNG tracks, in non-perturbed cells. Fig. 7D and E show that more tracks per cell can be seen for PolY2-mNG, compared to PolY1-mNG. However, frequently, replication forks did not colocalize with tracks of PolY2, indicating that the TLS polymerase is not a component of the replications forks, but only transiently recruited upon addition of DNA damage. Thus, although the copy number of PolY2 is higher than that of PolY1, possibly based on a higher basal transcription level even in the absence of SOS induction, there is also a clearly detectable heterogeneity in association with replication forks during non-perturbed conditions.

## Discussion

Since the discovery and biochemical characterization of DNA polymerases, it has become clear that replication forks must frequently deal with obstacles that not only slow down the progression of DNA replication, but often put a threat to the integrity of the entire assembled machinery (Bainbridge *et al*., 2021). Restart of stalled or even collapsed replication forks has been studied in great detail, especially at a biochemical and genetic level (Bainbridge *et al*., 2021; Torres *et al*., 2025; Windgassen *et al*., 2018). It has become clear that several different options exist for cells to overcome replication stress (Fig. 8), achieved by an armada of proteins that mediate very different avenues of DNA-based activities. Because biochemical and genetic approaches are population ensemble techniques, and single molecule biochemistry has not yet provided results on the investigation of entire forks plus some 30 proteins reported to play a role in restart, a clear view on the temporal activity of proteins at stalled forks is still missing.

**Fig. 8.**
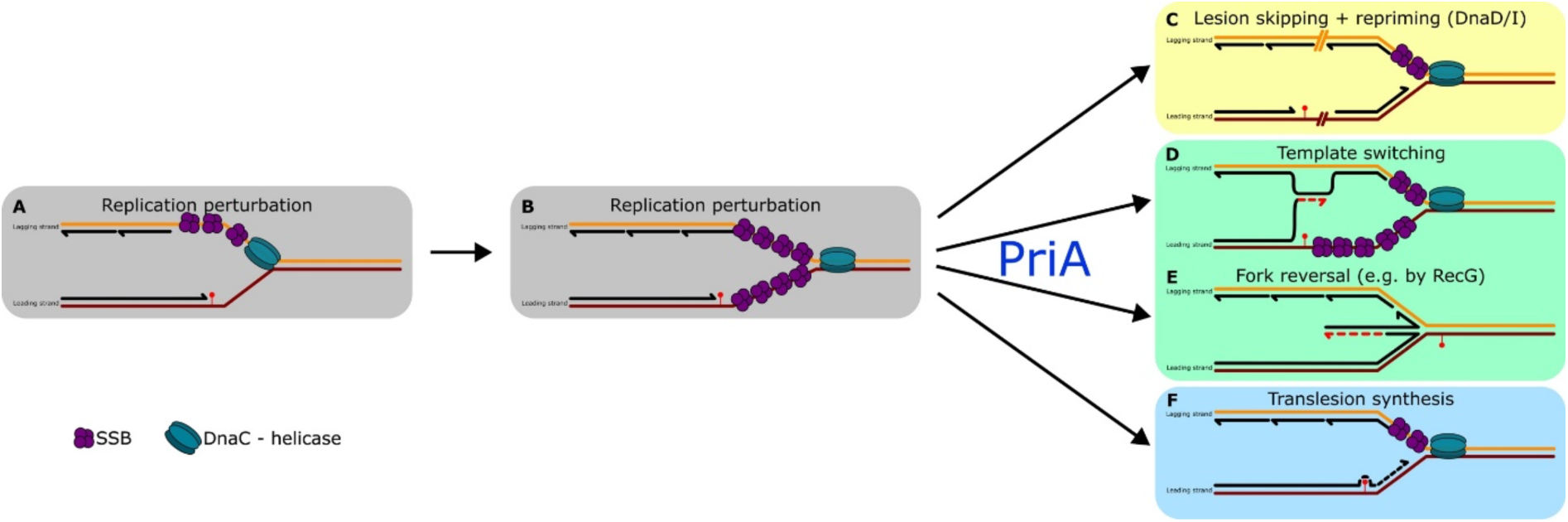
Model for major pathways to overcome replication obstacles on the leading strand. Adapted from (Torres *et al*., 2025).

We have employed epifluorescence microscopy and real-time single molecule tracking to study the localization and mobilities of key proteins in the resolution of replication stress in *B. subtilis* cells. A major conclusion of our studies is the heterogeneity of replication forks with respect to their composition. As opposed to an earlier study employing fluorescent proteins fused to restart proteins that were produced additionally to the endogenous proteins, we find that a functional mNG-PriA fusion expressed from the original gene locus is not generally pre-assembled at *B. subtilis* replication forks. While a subset (about 30%) of cells contained a PriA accumulation dependent on the C-terminus of SSB, most cells showed motion of PriA through the entire chromosome, with arrests at various positions. Thus, while some forks can rely on a proximity effect for PriA recruitment, most stalled forks depend on PriA targeting them by a transcription factor-like search for binding sites/structures. We show that pre-accumulation at the forks is also infrequent for primosome proteins DnaD and DnaI. Thus, like PriA, DnaD and DnaI and not general components of non-perturbed replication forks.

Interestingly, the number of mNG-PriA foci strongly increased after induction of replication stress. A large majority of cells showed 2 or more foci after stalling of PolC or inhibition of fork progression by MMC. Under the growth condition employed in this study, between 2 to 4 replication forks are present (Sharpe *et al*, 1998). We have used a concentration of MMC (added for 20 minutes) that is survived by about 75% of all cells (Kidane & Graumann, 2005). It is possible that this dose of MMC does not lead to an effect the progression of all replication forks, but it is reasonable to assume that a large majority of replication machineries witness replication stress. Our finding that PriA accumulated as foci in about 90% of all cells suggests that the major restart protein is indeed recruited to most stalled forks, no matter which pathway ensues to overcome the replication barrier (Fig. 8).

A second major conclusion is our finding that SSB shows a dramatic change in its single molecule motion following replication stress. In non-perturbed cells, about two-thirds of all molecules show free diffusion, while a third is in a slow mobile/static state consistent with binding to single strand DNA at replication forks. Stalling of PolC inverts this distribution, about 80% of SSB molecules change into a static mode, or more than 60% after treatment with MMC. This implies that despite the stalling polymerase progression, DnaC helicase continues to produce ssDNA in most cells. This would imply that the progression of DNA helicase does not seem to be as tightly coupled to polymerase activity in *Bacillus* compared with eukaryotic cells (Kim *et al*, 1996). Alternatively, DNA helicases such as PriA might unwind newly polymerized strands, in order to provide substrate for RecA-dependent recombination (template switching, Fig. 8D). In both scenarios, recruitment of SSB to forks under replication stress appears to be a first line of response, triggering (possibly alternative) downstream events.

SSB has been shown to be an important player in the response to replication stress and acts as an interaction or recruitment hub (Costes *et al*, 2010; Shinn *et al*, 2019). Our single molecule data showing that SSB is massively accumulating at stalled forks is not accompanied by a drastic switch from mobile to statically positioned molecules for PriA, DnaD or DnaI. We propose that rather than mediating a proximity effect, SSB acts in orienting the activity of PriA, and other restart proteins. Indeed, *in vitro*, *E. coli* SSB enhances the ability of PriA to discriminate between fork substrates as much as 140-fold. This is due to a significant increase in the catalytic efficiency of the helicase induced by SSB (Tan & Bianco, 2021). Our data show that PriA is generally less present in a static mode in the absence of the C-terminus of SSB, be it during non-perturbed growth, or in conditions of replication stress. Thus, our findings are in support of the idea that SSB plays an important role in enhancing PriA action at stalled forks, but not via establishing of a proximity effect.

DnaD and DnaI did not reveal accumulation at forks upon inhibition of PolC, and only to some degree after MMC treatment. These findings suggest lesion skipping and repriming is not a major route for dealing with obstacles on the leading strand (Fig. 8b). Stalling of DNA polymerase does not appear to invoke the loading of a new helicase hexamer at all, but may rely on a stabilization of arising ssDNA regions and a final catching-up of the polymerase if enzymatic inhibition is released. Under conditions when the progression of forks is hindered by chemical modification of bases or DNA interstrand crosslinks by MMC, some degree of new loading of DnaC appears to occur. PriA was found to accumulate in a vast majority of cells and at most replication forks, showing that this is a crucial step in replication restart that does not necessarily lead to the recruitment of DnaD, DnaI and likely also DnaB.

Heterogeneity of replication forks regarding association of restart proteins also extends to TLS polymerases. We show that PolY1 is present in exceedingly low copy number, such that in many cells, not a single molecule is present. Even in cells having multiple replication forks due to overlapping rounds of replication, as in our current study, a large number of cells contained few molecules, such that during replication stress, the TLS polymerase is not available at every fork. These findings do not agree with an earlier report that PolY1 is a general constituent of *B. subtilis* replication forks (Marrin *et al*., 2024). Also for PolY2, we found very low copy numbers in non-perturbed cells, revealing a limitation for the presence at replication forks, which must be found by chromosome scanning, rather than pre-association. Thus, the pathway of translesion synthesis (Fig. 8e) is highly limited concerning the availability of enzyme copies to perform the task. Surprisingly, PolY2 was present at higher copy number than PolY1, although it had been assumed that the gene encoding PolY2 (*yqjW*) is not expressed under normal conditions, but only in response to DNA damage via the SOS pathway (Au *et al*, 2005; Sung *et al*., 2003). Thus, PolY1 and PolY2 are examples for the importance of testing actual protein numbers in the cell, rather than deducing abundance from transcription data.

In conclusion, our data suggest that replication stress induced by the stalling of the leading (and lagging) strand DNA polymerase in *B. subtilis*, and by MMC-induced base modifications and interstrand crosslinks is largely overcome employing the pathway of fork remodelling and DNA synthesis, e.g. by fork reversal or template switching (Fig. 8c and d). This is supported by our observation that in contrast to the lack if the SSB interaction hub via the C-terminus, which leads to less strong (static) engagement of PriA after induction of replication stress, absence of the major HR helicase RecG generates an increased and somewhat prolonged static engagement of PriA with the chromosome. In other words, if the RecG-mediated path in HR is blocked, PriA can not leave blocked forks as efficiently as in wild type cells, supporting that HR follows initial recruitment of PriA at many stalled forks. Alternatively, PriA may bind to D-loop structures whose presence is increased in growing cells due to the lack of RecG activity (Briggs *et al*, 2004; Sanchez *et al*, 2007). For both scenarios, our finding of increased dwell times for PriA in the absence of RecG emphasizes a strong connection between PriA action and the HR pathway activities. Thus, by visualizing key events at stalled replication forks at a single molecule level, our work highlights the necessity of replication forks to be flexible in assuming different pathways for restart, in case some restart systems are not available due to strong heterogeneity between cells. Heterogeneity likely contributes to asymmetric propagation of mutations or DNA rearrangements, similar to what was observed for the repair of methylated guanosine in *E. coli* (Uphoff *et al*, 2016).

## Author contributions

Robert Raatz performed all experiments concerning figures 1 to 6, and supplementary figures 1, and 3 to 5. Daniel Hammerl performed all experiments related to figure 7. Anna Kornyushenko has helped with experiments for Figures 2 and 4, and has performed experiments for Fig. S2. Peter Graumann conceived of the study, supervised all experiments and organized funding. Robert Raatz and Peter Graumann wrote the manuscript.

