## Supplementary figures and tables for "Hierarchy and heterogeneity in the recruitment of *B. subtilis* replication-restart proteins to stalled forks"

**Supplementary material**

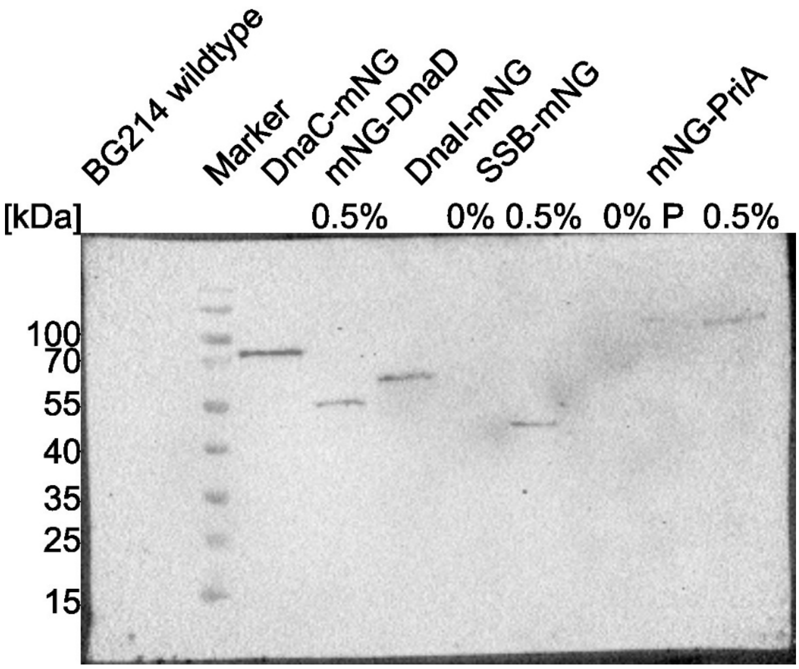

Fig. S1 Western blot showing mNeonGreen-fusions using anti-mNeonGreen (mNG) antibodies. Total cell extracts from exponentially growing cultures with xylose (0.5%) and without xylose (0%) in LB were used. Same amounts of protein were loaded in each lane. Expected band sizes for DnaC-mNeonGreen (77.3 kDa), mNG-DnaD (54.4 kDa), DnaI-mNG (62.9 kDa), SSB-mNG (45.5 kDa) and mNG-PriA(118.0 kDa) could be observed. mNeonGreen (26.9 kDa) was not observed. *Bacillus subtilis* BG214 WT was used as a control strain.

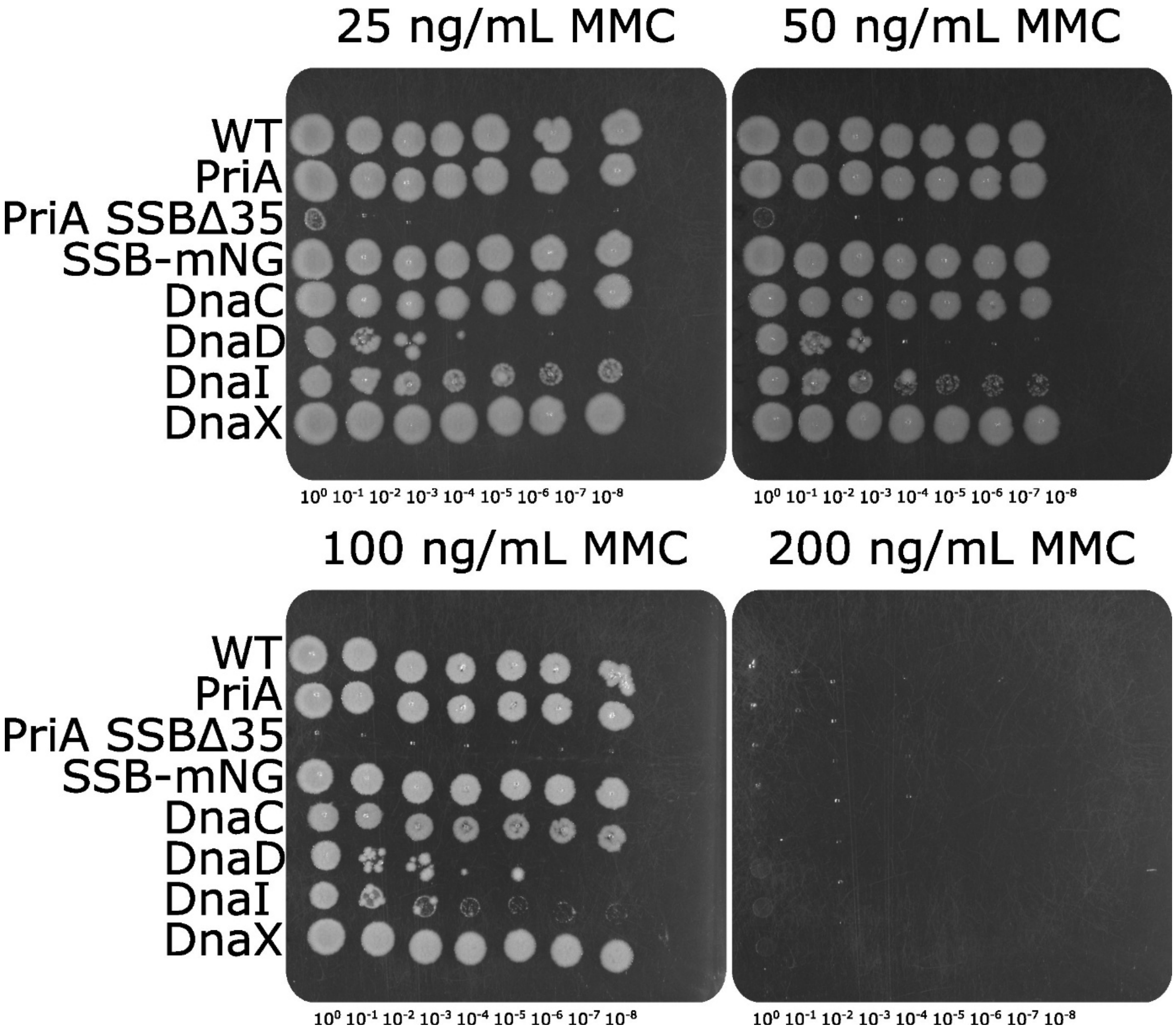

Figure S2: Susceptibility assay of the strains. LB-agar with different concentrations of MMC incubated at 30°C for approximately 24 h. mNG-PriA shows wild type-like susceptibility and SSBΔ35 shows expected sensitivity.

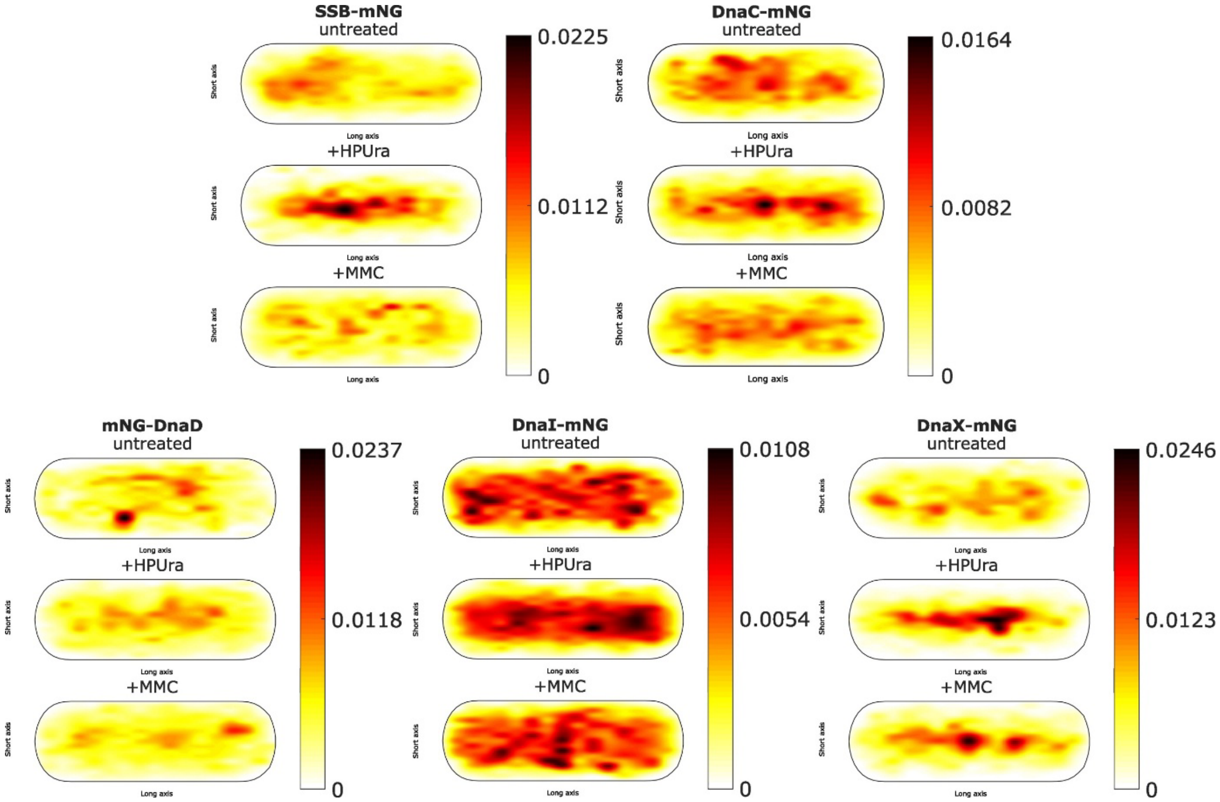

Fig. S3 Heat maps generated by SDA of SMTracker 2.0 for SSB-mNG, DnaC-mNG, mNG-DnaD, DnaI-mNG and DnaX-mNG. Proteins are shown untreated (upper panel), with HPUra (middle panel) and with MMC (lower panel). A scale for low (yellow) to high (black) indicates the probability.

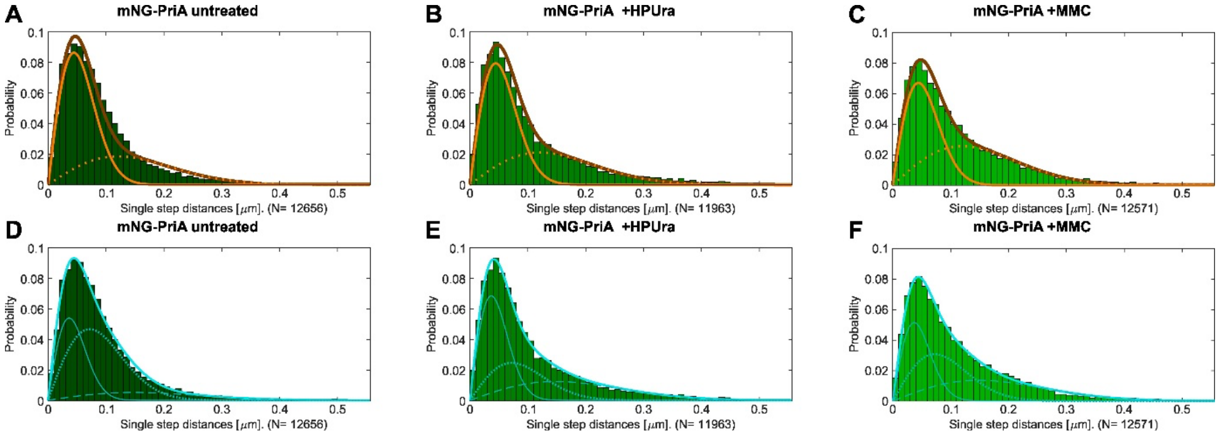

Fig S4 Jump distance analysis comparing 2 and 3 population fits. A,B and C show PriA in each condition assuming a double population fit. Single fits are shown in orange and the whole data coverage in brown. D, E and F show the same data assuming a triple fit. Single fits are shown in light blue and the whole data coverage in turquoise.

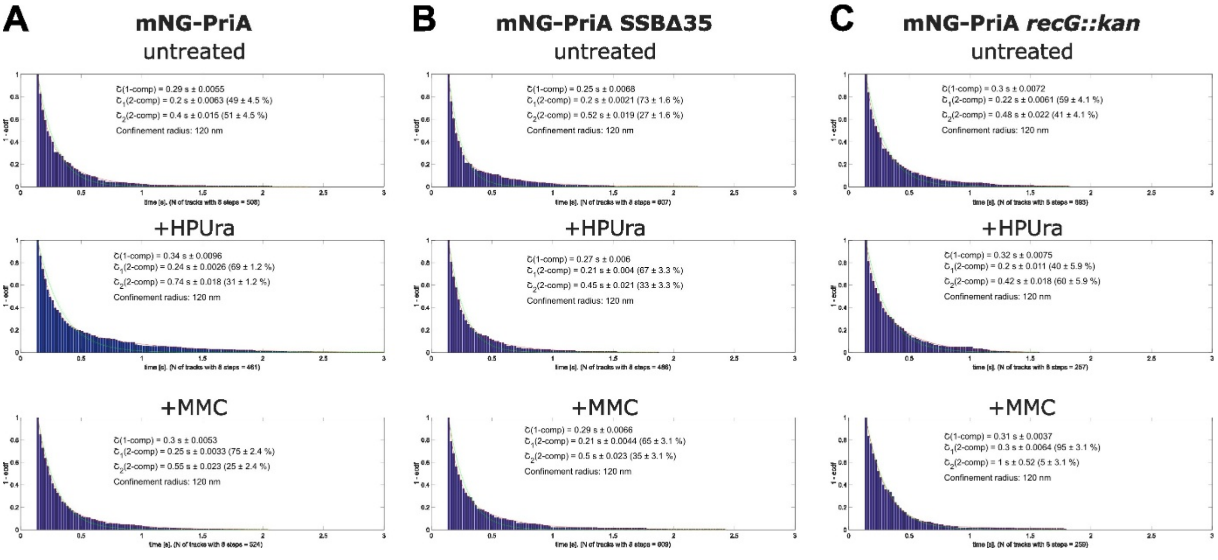

Fig. S5 Plots of dwell times for mNG-PriA in the mutant strains under each condition. Probability of PriA to stay within a radius of 120 nm for time t is depicted. A dwell-time cut-off of 8 frames was used. Dwell times were determined either as a 1- or 2-component fit using an exponential decay function.

**Table S1: Strains or plasmids used in this study**

| ***Strains or Plasmids*** | ***Relevant features*** | ***Reference or source*** |
|---|---|---|
| *E. coli* XL-1 Blue | | |
| *E. coli* BW25113 | | |
| *B. subtilis* BG214 | Wild type | Gift from Juan C. Alonso |
| PG4797 | *coaBC-mNG(*Δ3-5*)-priA* | This study |
| PG4798 | *coaBC-mNG(*Δ3-5*)-priA ssbA*Δ35 | This study |
| PG4799 | *amyE::Pxyl-ssbA-mNG-specR* | This study |
| PG4545 | *dnaC*-mNG-cmR *dnaX*-cfp-specR | (Hinrichs & Graumann, 2024) |
| PG4800 | *dnaX*-mNG-cmR | This study |
| PG4801 | *dnaI*-mNG-cmR | This study |
| PG4802 | cmR-Pxyl-mNG-*dnaD* | This study |
| 205 | *pPB105-ssbA*(Δ35)-2kb region; cmR | This study |
| 206 | *pPB105-coaBC-mNG(*Δ3-5*)-priA* 2kb region*;* cmR | This study |
| | *B. subtilis* 168 BKK15870 (Δ*recG::kan trpC2*) | (Koo *et al*, 2017) |
| PG4803 | *coaBC-mNG(*Δ3-5*)-priA* ΔrecG::kan | This study |
| PG3173 | *dnaX-cfp spec*$^R$ | (Lindow *et al*, 2002) |
| PG3730 | DH5α pSG1164-mVenus, expression Vector, AmpR, CmR | (Lucena *et al*, 2018) |
| 207 | pSG1164 dnaX-mNeonGreen, integration Vector, AmpR, CmR | This study |
| 208 | pSG1164 dnaI-mNeonGreen, integration Vector, Amp$^R$, Cm$^R$ | This study |
| 209 | pSG1193-*ssbA*-mNeonGreen, integration Vector, Amp$^R$, Spec$^R$ | This study |
| 210 | pHJDS-mNG-dnaD, integration vector, Amp$^R$, Cm$^R$ | This study |

Table S2: List of oligonucleotides

| | |
|---|---|
| **PriA - template parts** | |
| PriA_UP_fwd | GCATGCTGAATTCGTAATGAGGTTCctcatattgatgccgcagactgggc |
| PriA_UP_rev | GTTATCCTCCTCGCCCTTGCTCATgagcgctctccggtttgttttg |
| PriA_DOWN_fwd | AGGATCACTGCGGATCCACGGGTCCatgaattttgcagaagtcatcgttg |
| PriA_DOWN_rev | GCATAACCAAGCCTATGCCTACAGCtgaccatttgcggcgtcag |
| mNeonGreen(V2del)_fwd | ATGAGCAAGGGCGAGGAGGATAAC |
| mNeonGreen(V2del)_rev | GGACCCGTGGATCCGCAGTG |
| **PriA protospacer** | |
| PS_oligo1_mNGPriA_fwd | AAACatgacttctgcaaaattcatgagcgctctcG |
| PS_oligo2_mNGPriA_rev | AAAACgagagcgctcatgaattttgcagaagtcat |
| **ssbΔ35 - template parts** | |
| ssbΔ35_UP_fwd | GCATGCTGAATTCGTAATGAGGTTCgggcttcattcgtgctgagac |
| ssbΔ35_UP_rev | gcgataatcacattacccaaatggattatcattttggcc |
| ssbΔ35_DOWN_fwd | gataatccatttgggtaatgtgattatcgcctaaaatgaa |
| ssbΔ35_DOWN_rev | GCATAACCAAGCCTATGCCTACAGCtcaatccgccatccaatatttctg |
| **ssbΔ35 protospacer** | |
| PS_oligo1_ssbΔ35_fwd | AAACttgccaacgacggcaaaccgattgacatctG |
| PS_oligo2_ssbΔ35_rev | AAAACagatgtcaatcggtttgccgtcgttggcaa |
| **SSB-mNG (pSG1193)** | |
| pSG1193_ssb_rev | CAAGCTTGTCGCCCAGACTACCCGAgaatggaagatcatcatccgagatg |
| pSG1193_ssb_fwd | GATTCCTAGGATGGGTACCGGGCCCatgcttaaccgagttgtattagtcg |
| **DnaD (pHJDS-mNG)** | |
| pHJDS_dnaD_fwd | CTGCGGATCCACGGGTCCatgaaaaaacagcaatttattgatatgc |
| pHJDS_dnaD_rev | CTAGTGAATTCaacgcgtgtttgatcagttg |
| **DnaI (pSG1164-mNG)** | |
| pSG1164_dnaI_fwd | AGATTCCTAGGATGGGTACCttcagcatgttacagacttcc |
| pSG1164_dnaI_rev | AGGCCAGATAGGCCGGATCCtggatgtcggcggttttc |
| **DnaX (pSG1164-mNG)** | |
| pSG1164_dnaX_fwd | ATTCCTAGGATGGGTACCGAATTCcggaaagcccgaaaaaagaagc |
| pSG1164_dnaX_rev | TCCCAGGCCAGATAGGCCGGATCCgtcttttatttcaattaaatccgctccaac |
| ***recG::kan* LFH-primer from BKK** | |
| LFH_recGxkan_rev | gtgcagaatattcaccaaggctg |
| LFH_recGxkan_fwd | cggtaaagacattaaaaaaggcgac |
| Sequencing primer | |
| Seq_priA_rev | atcatcatataaggattcatatc |
| Seq_ssbΔ35_fwd | taatgtgattatcgcctaaaatgaaaaagaaagg |

| | |
|---|---|
| Seq_ssbΔ35_rev | cccaaatggattatcattttggcc |
| Seq_dnaD_rev | ccggaatttttggctgtgta |
| Seq_dnaI_fwd | aaacaacagtctcttatgaaaagcat |
| Seq_dnaX_fwd | gatttgccaaacctcacatcaatc |

**Table S3**

| | **PriA** | | | **PriA SSBΔ35** | | | **PriA *recG::kan*** | | |
|---|---|---|---|---|---|---|---|---|---|
| | **untreated** | **+HPUra** | **+MMC** | **untreated** | **+HPUra** | **+MMC** | **untreated** | **+HPUra** | **+MMC** |
| Dwell radius cut [nm] | 120 | 120 | 120 | 120 | 120 | 120 | 120 | 120 | 120 |
| Steps distribution free vs confined [%] | 60 / 40 | 62 / 38 | 72 / 28 | 75 / 25 | 77 / 23 | 79 / 21 | 66 / 34 | 53 / 47 | 57 / 43 |
| Average residence time [s] | 0.324 ± 0.013 s | 0.404 ± 0.025 s | 0.332 ± 0.016 s | 0.296 ± 0.014 s | 0.303 ± 0.015 s | 0.330 ± 0.020 s | 0.340 ± 0.013 s | 0.346 ± 0.018 s | 0.338 ± 0.018 s |
| $\tau$ (1-comp.) [s] | 0.29 ± 0.0053 s | 0.34 ± 0.0096 s | 0.3 ± 0.0054 s | 0.25 ± 0.0068 s | 0.27 ± 0.006 s | 0.29 ± 0.0067 s | 0.31 ± 0.0071 s | 0.32 ± 0.0075 s | 0.31 ± 0.0037 s |
| $\tau_1$ (2-comp.) [s] | 0.2 ± 0.0058 s | 0.24 ± 0.003 s | 0.25 ± 0.0035 s | 0.2 ± 0.0018 s | 0.21 ± 0.0039 s | 0.21 ± 0.0043 s | 0.22 ± 0.0057 s | 0.2 ± 0.011 s | 0.3 ± 0.0064 s |
| Fraction $\tau_1$ [%] | 47.9 ± 4.18 % | 69.3 ± 1.34 % | 75.8 ± 2.5 % | 72.6 ± 1.33 % | 67.3 ± 3.27 % | 64.7 ± 2.97 % | 58.5 ± 3.84 % | 39.7 ± 5.91 % | 95 ± 3.1 % |
| $\tau_2$ (2-comp.) [s] | 0.4 ± 0.014 s | 0.74 ± 0.021 s | 0.56 ± 0.027 s | 0.52 ± 0.015 s | 0.46 ± 0.021 s | 0.5 ± 0.022 s | 0.48 ± 0.02 s | 0.42 ± 0.018 s | 1 ± 0.52 s |
| Fraction $\tau_2$ [%] | 52.1 ± 4.18 % | 30.7 ± 1.34 % | 24.2 ± 2.5 % | 27.4 ± 1.33 % | 32.7 ± 3.27 % | 35.3 ± 2.97 % | 41.5 ± 3.84 % | 60.3 ± 5.91 % | 4.97 ± 3.1 % |
